# A functional genomics screen of human B-cell differentiation reveals convergent mechanisms of inherited childhood leukemia predisposition

**DOI:** 10.64898/2026.08.06.743305

**Authors:** Lara Wahlster, Anna-Lena Neehus, Andrew J. Lee, Soumyaa Mazumder, Pardiss Mehrzad, Susan Black, Luana Messa, Tanxin Liu, Charley Wang, Chen Weng, Alexis Caulier, Jensen Pak, Travis Fleming, Mateusz Antoszewski, Allison Zhang, Samuel A. Ha, Carmen Oleaga-Quintas, Adam J. de Smith, Vijay G. Sankaran

## Abstract

B-cell acute lymphoblastic leukemia (B-ALL) is the most common childhood cancer, yet the mechanisms by which inherited risk variants predispose to leukemia development remain poorly understood. A major challenge to studying these mechanisms has been the lack of model systems that faithfully capture the transient developmental states in which predisposition alleles are thought to act. Here, we establish a human B-cell differentiation platform from hematopoietic stem/progenitor cells that enables CRISPR-based engineering, recapitulates early B-cell lymphopoiesis, and enriches for rare developmental intermediates. By applying systematic perturbations with multiplexed single-cell transcriptomic profiling to mimic the effects of mutations in nine familial B-ALL predisposition genes, we decipher mechanisms by which B-cell development can be altered by such inherited variation to predispose to B-ALL. Through these studies, we identify convergent delays in B-cell differentiation at progenitor stages characterized by high-level RAG1/2 recombination activity. We propose that these delays at progenitor stages increase the likelihood that cells can undergo illegitimate RAG-mediated recombination to promote transformation, a finding consistent with similar rates of illegitimate RAG-associated genomic alterations in those with B-ALL associated with familial predisposition variants compared to sporadic cases.

## Introduction

Acute lymphoblastic leukemia (ALL) remains a leading disease-related cause of death in individuals under 20 years of age, with B-cell ALL (B-ALL) being the most prevalent subtype (1). While therapeutic advances have improved overall survival, many ALL survivors face a lifelong burden of treatment-related morbidity, and a subset of patients develop refractory disease with poor prognosis (2). To develop mechanism-based strategies for targeted prevention and treatment, a deeper understanding of the molecular origins of B-ALL is required. Substantial progress has been made in defining somatic driver mutations and their underlying mechanisms in B-ALL (3,4). Accumulating evidence demonstrates that inherited genetic variation plays an important and understudied role in predisposing individuals to B-ALL (5,6). Genome-wide association studies (GWAS) have identified more than 20 risk loci containing common variants linked to increased disease susceptibility, typically with low-to-moderate effect sizes in the range of ∼1.2 to 2.0 per risk allele, which collectively explain an approximately 10-fold variation in overall risk for acquiring B-ALL (7–11). In addition, genetic studies of familial B-ALL have identified rare germline variants that substantially increase disease risk. Although uncommon, these cases are particularly informative, revealing rare but highly penetrant variants in genes encoding hematopoietic transcription factors (TFs), DNA repair proteins, cell cycle regulators, cytokine signaling molecules, and protein deubiquitinates (5,12–20).

While previous functional studies have examined the role of specific familial B-ALL genes (12,21,22), a systematic analysis of these high-penetrance variants and how they might collectively impact cancer risk has been lacking, in part due to challenges in resolving and experimentally accessing the transient cellular states from which leukemic clones are thought to arise. The developmental origin of B-ALL has long been thought to lie within a window of vulnerability along the B-cell differentiation trajectory, generally ranging from the pro-B cell stage through early pre-B cells (23–27). More recently, a growing body of evidence suggests that the origins of B-ALL may be more heterogeneous than previously appreciated, with leukemic programs appearing to arise across a broader spectrum of early hematopoietic and B-lineage developmental states, depending on the molecular subtype (28). In line with this, comparative transcriptomic studies have shown that distinct B-ALL subtypes can exhibit transcriptional features consistent with differing degrees of lineage restriction, ranging from committed B-cell intermediates to more primitive progenitor-like states, depending on the genomic context of the disease (28–32). Notably, most current models of cellular origin rely on retrospective analyses of B-ALL clones, which can be influenced by clonal evolution, treatment exposure, and disease progression, complicating inference of the initiating cell state. In this context, variant-to-function mapping in single-cell multi-omic data from human hematopoietic progenitors has provided complementary evidence that common B-ALL risk variants are significantly enriched in regulatory elements active during key lineage transitions in early B-cell development, particularly across intermediates spanning the pro-B to naïve B-cell stages, with a peak in early pre-B cells (33). However, despite these insights, accessing and efficiently manipulating the relevant transient developmental states in primary human systems remains challenging, highlighting the need for scalable and tractable *in vitro* models that recapitulate early human B-cell development.

To overcome these limitations, we adapted and deeply characterized a robust *in vitro* human B-cell differentiation platform from hematopoietic stem and progenitor cells (HSPCs) that enables precise genetic perturbation. We employed this system to perform an arrayed functional genomic screen to gain systematic insights into the mechanisms by which these predisposing variants act by combining CRISPR/Cas9 genome editing in primary human HSPCs with multiplexed single-cell transcriptomic profiling across B-cell differentiation. Using this platform, we modeled pathogenic variants in nine genes implicated in familial B-ALL predisposition: *CDKN2A, ETV6, IKZF1, PAX5, USP9X, TCF3, TP53, SH2B3*, and *NBN* (12,14–16,18,19,22,34,35). Single-cell trajectories revealed that diverse loss-of-function (LOF) perturbations converge on distinct, stage-specific developmental bottlenecks during early B-cell lymphopoiesis - most notably at stages characterized by high-level RAG1/2 recombination activity - the precise mechanism that underlies genomic instability in B-ALL and causes driver lesions to arise. Critically, we show this is likely to be a mechanism that increases propensity for transformation, rather than intrinsic genome instability, by showing that the pattern of somatic structural alterations in over 1,400 childhood B-ALL cases does not differ in those with or without familial predisposition variants. Together, these findings define convergent mechanisms by which childhood B-ALL can arise and establish a scalable system for functional dissection of leukemia predisposition during human hematopoietic development.

## Results

### Differentiation of early B-cell progenitors and precursors from human hematopoietic stem and progenitor cells

To investigate how germline variants shape B-cell lymphopoiesis and predispose to the development of B-ALL, we utilized a robust *in vitro* culture system that recapitulates the full continuum of early human B-cell differentiation (**Figure 1A-B**). Previous studies have shown that co-culture of human HSPCs with mouse or human stromal cells, together with recombinant cytokine supplementation, can support B-lineage commitment (36–40). Building on these approaches, we adapted an *in vitro* stromal co-culture platform for B-cell differentiation from cord blood-derived CD34^+^ HSPCs (**Figure 1A**). The system employs a two-stage differentiation strategy consisting of a short expansion and lineage priming phase in serum-free medium supplemented with recombinant human fms-like tyrosine kinase 3 ligand (FLT3L), stem cell factor (SCF), interleukin 3 (IL-3), interleukin 6 (IL-6), and thrombopoietin (TPO), followed by long-term co-culture on MS-5 stromal monolayers supplemented with recombinant human interleukin 7 (IL-7) to support B-cell specification and maturation. To evaluate the developmental output of the system, we first assessed the morphology of CD19^+^ cells isolated after 5 weeks of co-culture on cytospin slides stained with May-Grünwald Giemsa. The *in vitro*-derived cells exhibited a small lymphoid morphology with a high nucleus-to-cytoplasm ratio, comparable to *bona fide* naïve B cells isolated from peripheral blood of healthy donors (**Figure 1C**). We next characterized the cellular composition of the culture by flow cytometry to resolve B-lineage differentiation dynamics over time. CD19^+^ B-lineage progenitors emerged as early as 14 days after initiation of the co-culture and increased progressively through week 5 of differentiation (**Figure 1D-E**). Cell surface staining with CD10 and CD20 further revealed distinct B-cell precursor stages within the culture, with a progressive increase in CD20^+^ cells over time consistent with ongoing maturation of the CD19^+^ compartment.

**Figure 1:**
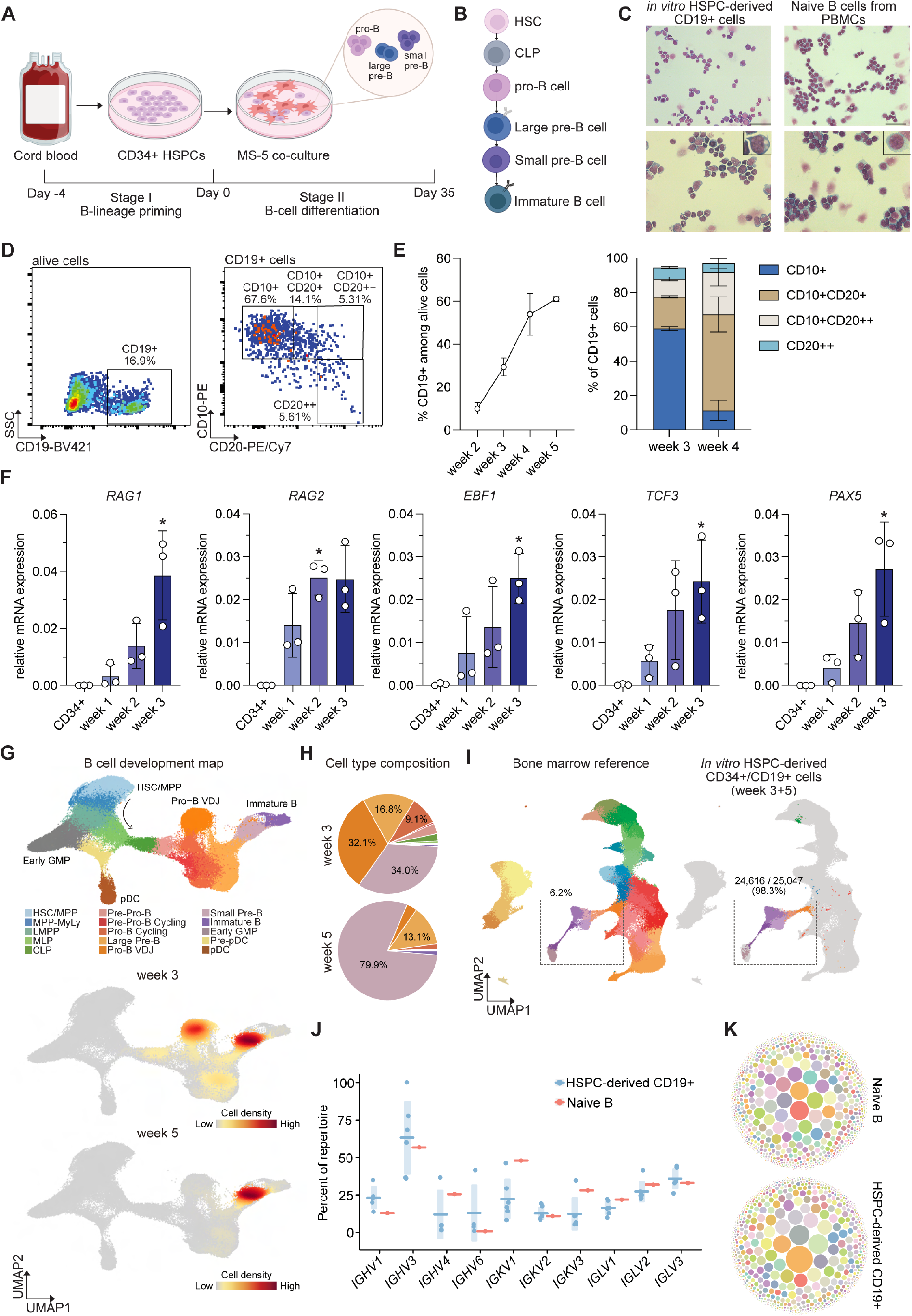
*In vitro* modeling of early human B cell lymphopoiesis. **(A)** Schematic overview of the two-stage differentiation strategy. **(B)** Schematic overview of human B lymphopoiesis. **(C)** Representative cytospin images with May-Grünwald-Giemsa staining of *in vitro*-derived CD19^+^ cells (left) and naive B cells isolated from healthy donor PBMCs (right). Scale bar: 50 µm. **(D)** Representative flow cytometry staining for CD19, CD10 and CD20 after 3 weeks of *in vitro* differentiation cultures. **(E)** Longitudinal quantification of CD19^+^ cells and theirCD10/CD20 expression profile from one representative donor. Data are shown as mean ± SD. **(F)** Quantitative RT-PCR analysis for key B cell marker genes relative to *GAPDH* in unsorted differentiation cultures over time (*n*= 3). Data are shown as mean ± SD. Statistical significance was assessed using Friedman test with Dunn’s multiple-comparisons test compared to CD34^+^. *P < 0.05. **(G)** UMAP of scRNA-seq data obtained from HSPC-derived cells sorted for CD34^+^ and CD19^+^ after 3 and 5 weeks of differentiation culture, colored by cell type, together with UMAP plots showing the density of cells at 3 and 5 weeks of culture along the B cell developmental trajectory. **(H)** Pie charts showing the proportion of scRNA-seq cells by cell type. Colored as in **(G). (I)** Projection of HSPC-derived scRNA-seq cells onto a human bone marrow reference map, highlighting the proportion of cells within B cell progenitor states. **(J-K)** B-cell receptor repertoire analyses from HSPC-derived CD19^+^ B cells (*n* = 5) and naive B cells isolated from healthy donor PBMCs (*n* = 1). Usage of different *IGH, IGK* and *IGL* gene elements **(J)** and bubble plots depicting the clonal diversity of HSPC-derived CD19^+^ cells compared with naïve B cells **(K)**.

To validate activation of canonical B-cell transcriptional programs in this culture system, we performedvquantitative RT-PCR on unsorted differentiating cultures and observed temporal induction of key B-lineagevtranscription factors (*EBF1, TCF3* and *PAX5*) alongside progressive upregulation of the genes encoding enzymes RAG1 and RAG2, which are critical for V(D)J recombination (**Figure 1F**). To resolve the full developmental architecture of the system at single-cell resolution, we performed single-cell RNA sequencing (scRNA-seq) on co-cultures at 3 weeks and 5 weeks of differentiation following enrichment for CD34^+^ and CD19^+^ fractions. This analysis revealed a continuum of early B-cell developmental states, capturing lymphoid-primed multipotent progenitors (LMPPs), common lymphoid progenitors (CLPs), pre-pro B cells, pro-B cells, and pre-B cells (**Figure 1G-H**). These HSPC-derived B cell progenitors closely recapitulated native human bone marrow counterparts, exhibiting comparable expression of key B lineage drivers and markers, as well as highly similar global transcriptional profiles (**Supplementary Figure 1A-B**) (41). Comparison between week 3 and week 5 cultures highlighted the progressive developmental maturation within the system, consistent with our flow cytometry data (**Figure 1D-E**). Whereas week 3 cultures maintained a broader spectrum of early B-lineage intermediates, including CLPs, pre-pro-B, and pro-B cells, week 5 cultures were predominantly composed of small pre-B cells, indicating continued and coordinated differentiation over time (**Figure 1G-H**). Benchmarking this co-culture system against primary human bone marrow aspirates (42) demonstrated that the system faithfully recapitulates early B-lineage hierarchies, while strongly enriching for rare transitional populations, including pro-and pre-B cells that are generally poorly captured in primary tissue samples (**Figure 1I**). Notably, the frequency of these highly transient developmental states increased from 6.3% of total bone marrow mononuclear cells to 98.3% of the cells in our *in vitro* co-cultures (**Figure 1I)**. Finally, B-cell receptor (BCR) profiling confirmed progressive immunoglobulin gene rearrangement with diverse *IGHV, IGKV*, and *IGLV* gene segment usage, consistent with highly polyclonal B-cells whose BCR diversity closely resembled that of primary human naive B cells (**Figure 1J-K, Supplementary Figure 1C-D**). Collectively, these results demonstrate that this co-culture system of human HSPCs yields a semi-synchronous continuum of early B-cell development, capturing key differentiation stages relevant to the pathogenesis of B-ALL (43).

### Efficient CRISPR/Cas9 perturbation of familial B-ALL predisposition genes defines their effects in early B-cell differentiation

To demonstrate the utility of this platform for systematic functional interrogation of leukemia predisposition variants, we selected nine genes implicated in familial and/or syndromic forms of B-ALL, including *CDKN2A, ETV6, IKZF1, NBN, PAX5, SH2B3, TCF3, TP53*, and *USP9X* (**Figure 2A, Supplementary Figure 2**) to conduct a single-cell arrayed perturbation screen. These genes have been associated with B-ALL predisposition through germline hypomorphic or loss-of-function (LOF) heterozygous variants, with the exception of *SH2B3*, which has primarily been reported in the context of homozygous inheritance (12–19,22,34,35,44). To maximize the fidelity and robustness of genetic modeling, we designed multiple candidate single guide RNAs (sgRNAs) targeting coding regions in close proximity to reported pathogenic germline variants and/or within critical functional domains (**Supplementary Figure 2**) and selected two sgRNAs per target with the highest editing efficiency for downstream analyses (**Supplementary Figure 3A**). Two sgRNAs targeting the AAVS1 safe-harbor locus served as negative controls. Electroporation of human HSPCs with Cas9 ribonucleoprotein complexes resulted in high levels of locus-specific editing across all target genes. Importantly, editing efficiency remained stable throughout differentiation for the majority of sgRNAs tested, both five days after editing and following three weeks of co-culture (**Figure 2B**), enabling the interrogation of gene function during the transient and developmentally sensitive phases in which B-ALL predisposition alleles are likely to act. Sequence analysis demonstrated that editing predominantly generated frameshift alleles predicted to disrupt protein function and/or expression, thereby mimicking the pathogenic effect of the known B-ALL predisposition alleles (**Supplementary Figure 3B**). One notable exception was observed at the *IKZF1* locus, where one sgRNA primarily generated predicted LOF alleles, whereas a second sgRNA reproducibly produced a six-nucleotide in-frame deletion impacting the N-terminal zinc finger 4 domain of IKAROS (**Supplementary Figure 3C**) in which missense variants have been reported to underlie early-onset combined immunodeficiency via a dominant negative (DN) effect (45).

**Figure 2:**
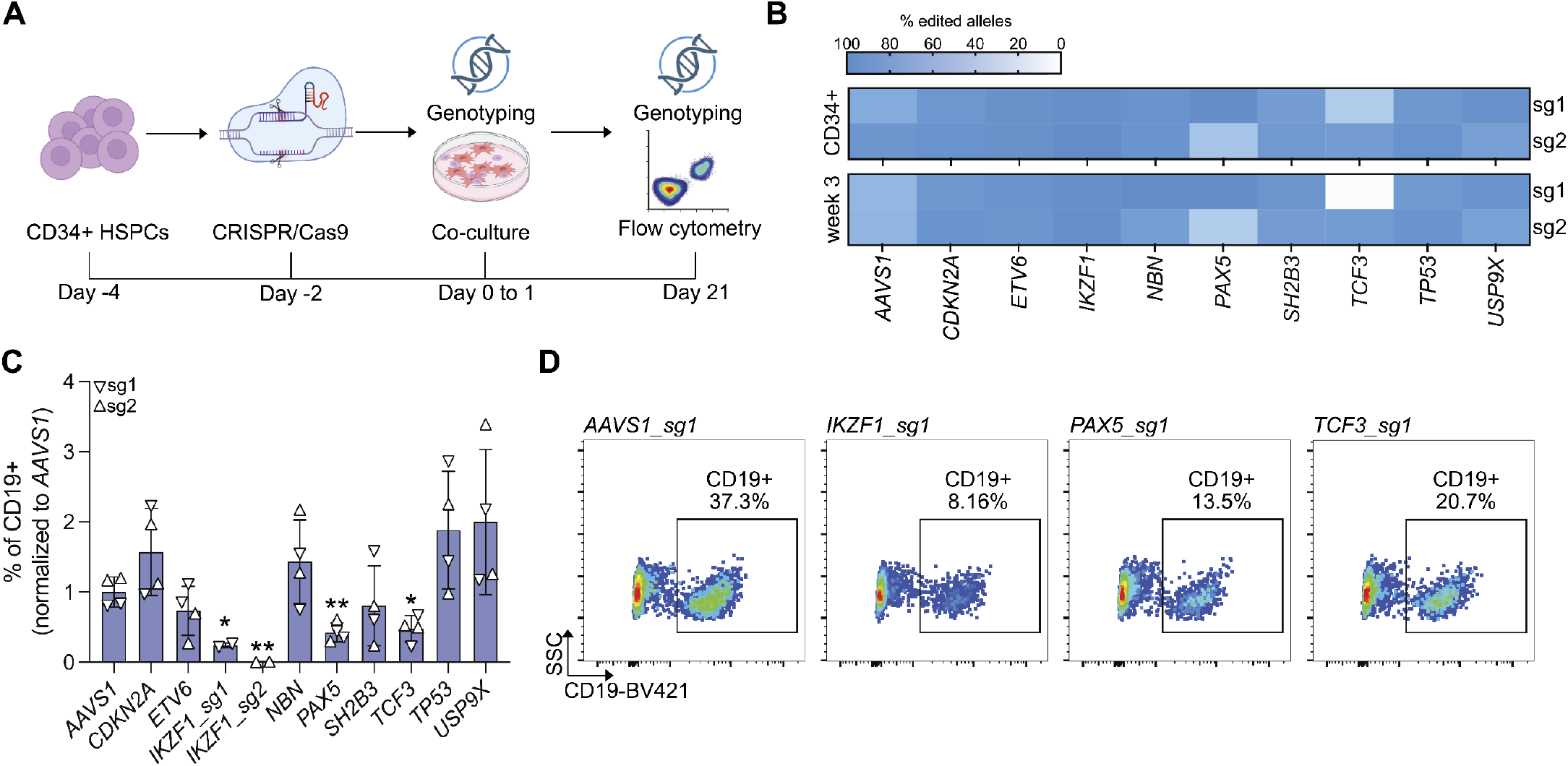
CRISPR/Cas9 modeling of B-ALL predisposition loci. **(A)** Schematic overview of the experimental design for CRISPR/Cas9-editing of human CD34^+^ HSPCs, followed by genotyping, MS-5 co-culture and flow cytometric analysis. **(B)** Editing efficiencies across nine targeted B-ALL risk loci in CD34^+^ HSPCs and at week 3 of co-culture across two independent sgRNAs (sg1, sg2) per target gene. **(C)** Flow cytometric quantification of the relative frequency of committed CD19^+^ B-lineage cells harvested at week 3 of co-cultures, with values normalized against the mean of the AAVS1 control. Data are shown as mean ± SD across two biological replicates with two guides per target. Statistical significance was assessed using one-sample t-tests. *P < 0.05, **P < 0.01. **(D)** Representative flow cytometry staining for CD19 at week 3 of co-culture across indicated sgRNAs.

Flow cytometric analysis after three weeks of differentiation revealed gene-specific effects on early B-cell development. Disruption of the key B-lineage transcription factors *IKZF1*, PAX5, and *TCF3* resulted in marked reduction in CD19^+^ cells, consistent with impaired B-cell commitment and maturation (**Figure 2C-D**; **Supplementary Figure 3D**). The most pronounced phenotype was observed with the *IKZF1* sgRNA generating the six-nucleotide in-frame deletion, suggesting that distinct classes of *IKZF1* mutations exert differential effects on B-lymphopoiesis, mirroring phenotypic variations observed in patient cohorts (**Supplementary Figure 3D**) (46). In contrast, CRISPR/Cas9-mediated disruption of the remaining six predisposition genes did not significantly alter overall output of CD19^+^ cell under these baseline culture conditions, although this marker cannot discriminate if alterations in B-cell development are arising, given its continuous expression over much of B-lymphopoiesis (**Figure 2C**). Together, these findings demonstrate that perturbation of familial B-ALL genes exerts diverse effects on early lymphopoiesis, ranging from overt blocks in B-cell production to phenotypes that remain superficially unperturbed at the bulk level. This variation in total cellular output highlights the limitations of conventional bulk phenotyping and suggests that predisposition-associated defects may manifest as alterations in developmental transitions, rather than changes in overall lineage output.

**Figure 3:**
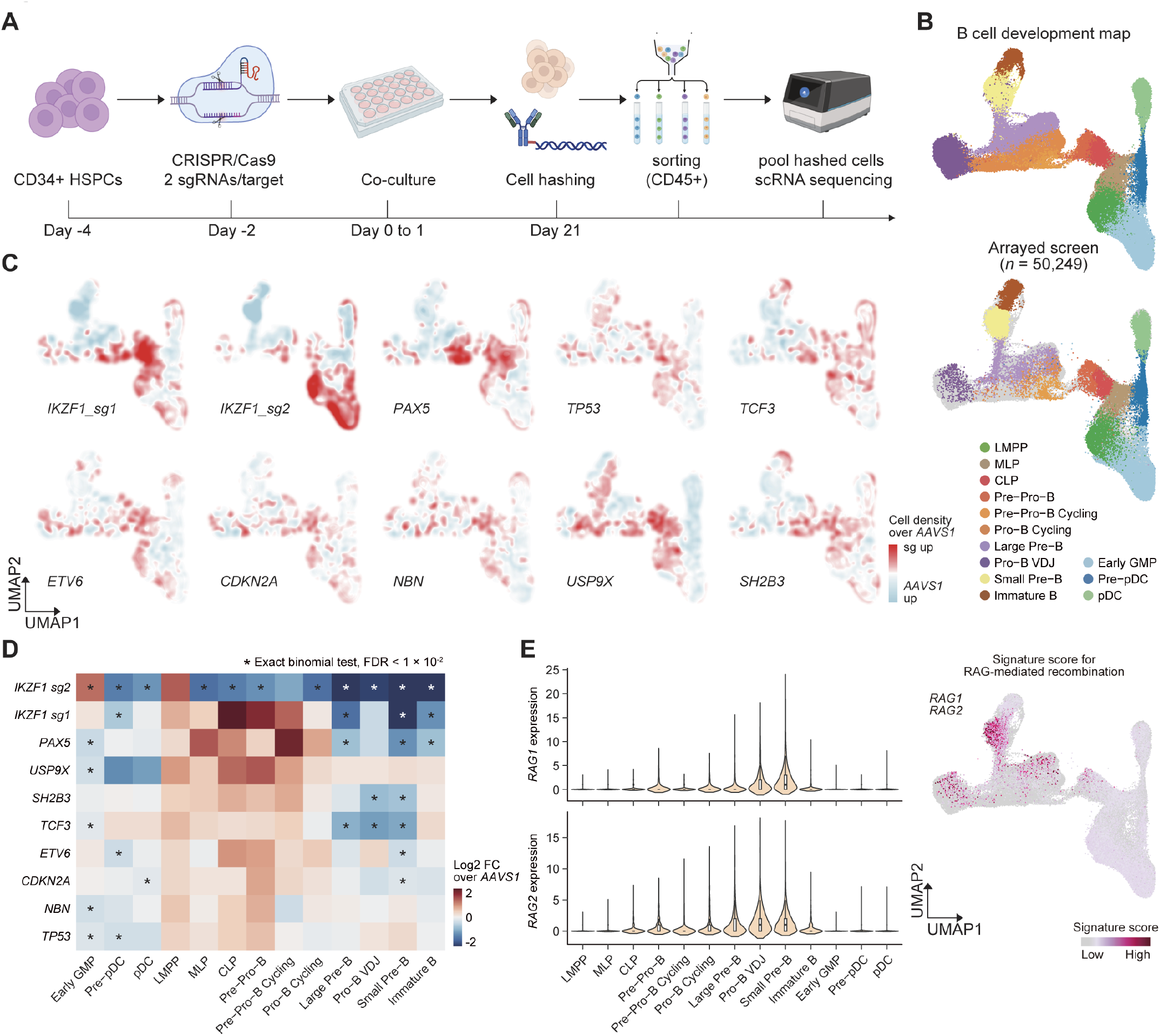
A multiplexed single-cell CRISPR screening strategy for functional evaluation of B-ALL risk mutations. **(A)** Schematic overview of the experimental design for the hashed arrayed screen using human CD34^+^ HSPCs. **(B)** UMAP showing the arrayed screen scRNA-seq cell projection onto a human B cell development reference map. **(C)** Cell density profiles across gene perturbations showing the relative cell density ratio relative to *AAVS1* control-edited cells. **(D)** Quantitative enrichment and depletion analysis across targeted B-ALL predisposition loci. The heatmap displays the log2 fold enrichment or depletion relative to the *AAVS1* control. Statistical significance was assessed using a one-sided exact binomial test, with false discovery rate (FDR) correction for multiple testing. *FDR < 0.01. **(E)** Violin plots showing RAG1 and RAG2 expression across different stages of B cell lymphopoiesis (left) and a UMAP plot of transcriptional signature scores for V(D)J recombination represented by *RAG1* and *RAG2* expression (right).

### B-ALL risk mutations stall B cell development at distinct differentiation stages

To systematically delineate the functional consequences of B-ALL predisposition variants on early B-cell development, we implemented a multiplexed single-cell CRISPR screening strategy designed to resolve both robust and subtle perturbation effects across developmental trajectories. To enable multiplexed scRNA-seq across multiple gene edits in parallel, we performed an arrayed perturbation approach, combined with cell hashing using barcoded oligo-conjugated antibodies to conduct this CRISPR perturbation-based functional screen (**Figure 3A**) (47). Using this framework, CD34^+^ human HSPCs were edited with CRISPR/Cas9 using two sgRNAs per target across the nine B-ALL predisposition genes. Following three weeks of *in vitro* B-lineage differentiation to ensure robust sampling of transient early B-cell progenitor populations, the distinct experimental conditions were labeled with unique oligo-conjugated surface antibodies to enable cell hashing (**Figure 3A**). To ensure that early differentiation bottlenecks and lineage diversion events were not excluded a priori, we sorted for CD45^+^ cells to capture the full spectrum of hematopoietic and lymphoid states emerging during *in vitro* differentiation. In total, we obtained high-resolution transcriptomic readouts for 50,248 cells across all targeted perturbations. To resolve precise cellular identities, these single cell profiles were projected onto a reference map of normal human B-cell development obtained from the bone marrow (43). Our platform successfully recovered cells spanning the entire physiological continuum of early human lymphopoiesis, continuously capturing trajectories from common lymphoid progenitors (CLPs) through pro-B and pre-B cell intermediates to immature B cells (**Figure 3B**; **Supplementary Figure 4A-B**).

**Figure 4:**
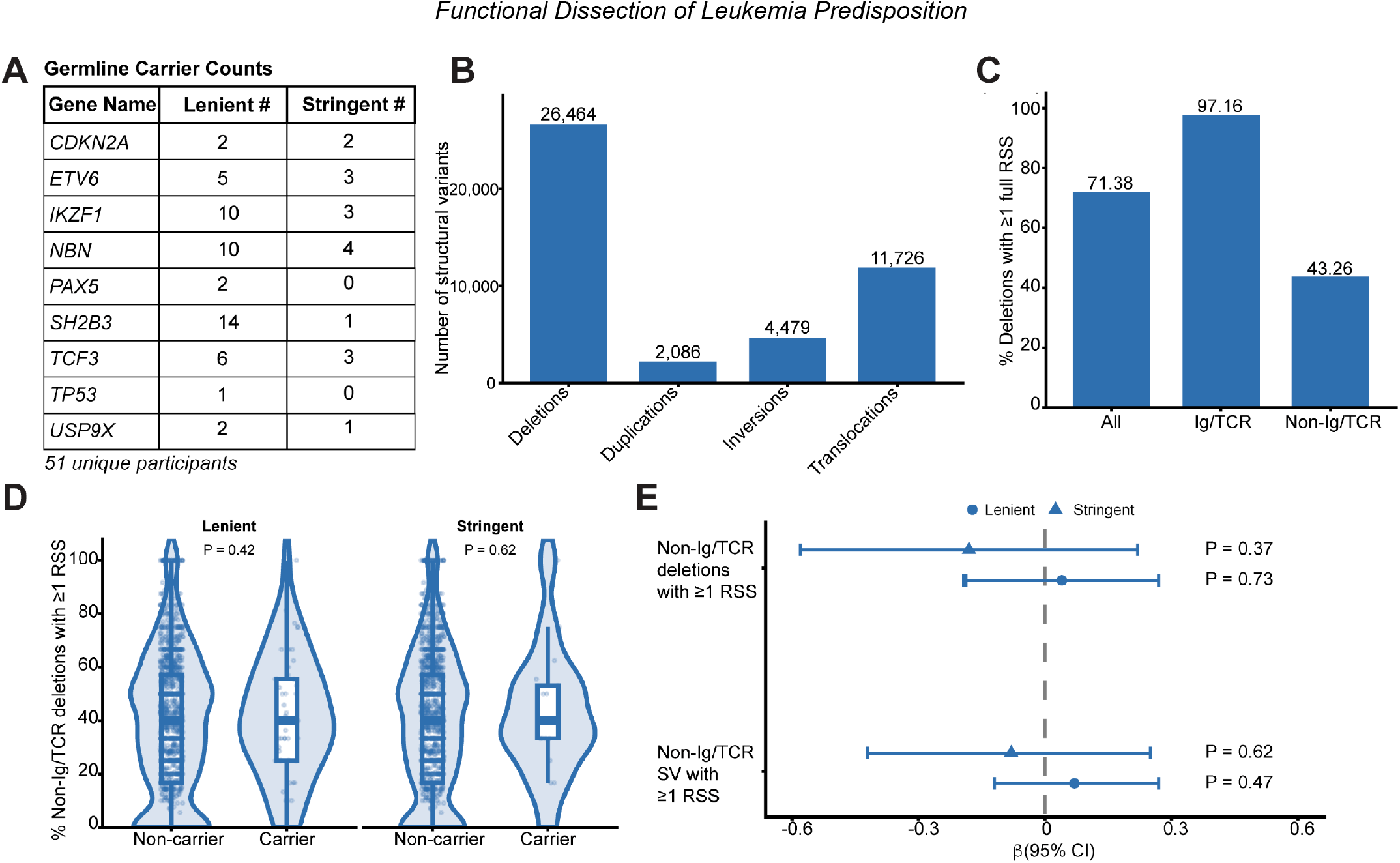
RAG-associated structural variation in B-ALL cases with and without germline predisposition variants. **(A)** Gene-specific numbers of patients harboring pathogenic or likely pathogenic germline variants in nine B-ALL predisposition genes under lenient and stringent variant-classification criteria. Gene-specific counts are not mutually exclusive. **(B)** Numbers of somatic structural variants detected using BRASS across 1,491 pediatric B-ALL cases. **(C)** Percentage of deletions containing at least one full-length recombination signal sequence motif at either breakpoint across all deletions, deletions involving immunoglobulin or T-cell receptor loci, and deletions outside Ig/TCR loci. **(D)** Distribution of the proportion of non-Ig/TCR deletions containing at least one full-length RSS motif among patients with and without P/LP germline variants under lenient and stringent classification criteria. Violin plots show the distribution across patients, and box plots indicate the median and interquartile range. Patients without non-Ig/TCR deletions were excluded because the proportion was undefined. Statistical significance was assessed using a two-sided Wilcoxon rank-sum test. **(E)** Multivariable negative binomial regression analysis of the association between germline P/LP variant carrier status and the number of RSS-associated non-Ig/TCR deletions or structural variants. Models were adjusted for age at diagnosis and molecular subtype and were evaluated separately under lenient and stringent variant criteria. Points represent regression coefficients, and horizontal lines indicate 95% confidence intervals.

To systematically delineate how individual germline risk variants shape cell fate, we performed single-cell density mapping and guide-specific enrichment analyses (**Figure 3C-D**). This approach revealed highly reproducible, perturbation-specific differentiation bottlenecks across the targeted B-ALL predisposition genes, with the exact developmental perturbation varying by genetic locus (**Figure 3C-D**). Disruption of the core B-lineage transcription factor PAX5 resulted in a strong block at early B-lineage commitment. Single-cell density profiles showed a marked accumulation of cells at the CLP and Pre-Pro-B cell stages, accompanied by a pronounced depletion of downstream populations. In contrast, the two *IKZF1* perturbations resolved distinct functional modalities consistent with their predicted LOF and DN effects, respectively (**Figure 3C-D**). The LOF sgRNA (IKZF1_sg1) resulted in an accumulation of cells within the early lymphoid compartment, including CLP and Pre-Pro-B cell stages, consistent with a partial developmental block, similar to the effects of disrupting *PAX5*. By comparison, the DN sgRNA (IKZF1_sg2) induced a more severe collapse of the B lineage trajectory, characterized by a near-complete depletion of committed early B-cell progenitors, alongside a reciprocal accumulation in more primitive multipotent progenitor populations like LMPPs (**Figure 3C-D**). Beyond these strong arrest phenotypes, our multiplexed scRNA-seq platform also resolved more subtle, stage-restricted vulnerabilities among additional predisposition genes, underscoring the value of this high-resolution single-cell arrayed perturbation screen. Perturbations in *CDKN2A, ETV6*, and *SH2B3* largely preserved overall CD19^+^ B-cell output from our *in vitro* cultures, as shown by flow cytometry (**Figure 2C**), but exhibited a shared depletion signature around the small pre-B cell developmental state, indicating that genetic loss of these loci impairs progression through early B-lymphopoiesis (**Figure 3D**). Notably, the stages at which these bottlenecks accumulate correspond to periods of peak RAG1 and RAG2 expression during normal B-cell differentiation (**Figure 3E**), when V(D)J recombination is most active and the probability of off-target RAG-mediated cleavage at non-immunoglobulin loci is highest.

To investigate whether the observed differentiation bottlenecks converge on shared transcriptional programs or instead reflect gene-specific regulatory mechanisms, we assessed downstream transcriptional consequences for each perturbation. We reasoned that conventional global differential expression analyses would be confounded by the substantial shifts in cellular composition across B-ALL predisposition alleles (**Figure 3C-D**). We therefore restricted transcriptional comparisons to the specific developmental stages in which each perturbation-induced alteration was initially localized. Gene set enrichment analysis across the individual perturbation-stage comparisons revealed 35 significantly altered biological pathways (false discovery rate <0.01), including pathways involved in cell cycle regulation, cellular respiration, B cell development, and immune function (**Supplementary Figure 4C**). However, these pathway alterations were highly perturbation-specific, and we did not identify a consistent transcriptional program that was shared across all perturbations. Instead, each perturbation exhibited distinct transcriptional signatures, suggesting that these B-ALL predisposition alleles act on B-cell development through gene-specific regulatory programs that converge functionally on impaired B-cell maturation, but not necessarily at the molecular level. Together, these results demonstrate that our arrayed perturbation screen enables high-resolution dissection of genetic effects during early human B-cell development. Our findings reveal that high-penetrance germline risk variants converge functionally on impaired progression through early human B lymphopoiesis, with bottlenecks occurring at developmental stages characterized by elevated RAG1 and RAG2 expression and activity (**Figure 3E**).

### B-ALL predisposition alleles increase abundance of RAG-dependent structural variants

Structural variants (SV) that act as drivers in B-ALL have previously been shown to arise predominantly through off-target RAG-mediated illegitimate V(D)J recombination (48– 50). While our perturbation screen results suggest that predisposition variants may act by prolonging residence within developmental states characterized by high RAG1/RAG2 expression, thereby increasing the probability of acquiring illegitimate RAG-mediated driver rearrangements, we hypothesized that once such initiating events have occurred, the subsequent somatic mutational landscape in transformed clones should be comparable between carriers and non-carriers of predisposition alleles. To directly address this hypothesis, we analyzed whole-genome sequencing data from 1,491 pediatric B-ALL patients enrolled in the Children’s Oncology Group Molecular Profiling to Predict Response to Treatment (MP2PRT) cohort (51). This dataset includes paired germline and tumour samples, allowing comprehensive assessment of both inherited variation and somatic structural rearrangements. We first probed for pathogenic or likely pathogenic (P/LP) germline variants in the nine B-ALL predisposition genes that we previously modelled: *CDKN2A, ETV6, IKZF1, NBN, PAX5, SH2B3, TCF3, TP53*, and *USP9X*. Using lenient and stringent criteria, we identified 40 variants in 51 patients and 16 variants in 17 patients, respectively (**Figure 4A, Supplementary Table 4**). The most recurrently altered genes across both thresholds were *IKZF1, NBN*, and *TCF3*. To evaluate the somatic SV landscape in these individuals, we analyzed 44,755 structural variants detected using BRASS across the full cohort, including 26,464 deletions, 2,086 duplications, 4,479 inversions, and 11,726 translocations (**Figure 4B**). We next assessed whether these SVs harbored sequence motifs consistent with RAG-mediated cleavage. Motif enrichment analysis at SV breakpoints identified full recombination signal sequences (RSS) by detecting heptamer and nonamer motifs separated by canonical 12-or 23-base-pair spacers. Among all SV types, deletions were most strongly associated with RSS motifs. Of the 26,464 deletions detected genome-wide, 71.38% (18,891) harbored the full RSS motif in at least one breakpoint sequence, with higher enrichment in immunoglobulin and T-cell receptor (Ig/TCR) loci (13,416 of 13,808 deletions, 97.16%), as expected (**Figure 4C**) (52). Importantly, 43.26% of deletions (5,475 of 12,656) outside Ig/TCR regions (non-Ig/TCR) showed RSS enrichment. To determine whether germline predisposition is associated with susceptibility to RAG-mediated SVs, we then stratified patients by P/LP variant status. We find that B-ALL cases harboring P/LP germline variants exhibited SVs with hallmarks of RAG-mediated recombination at frequencies comparable to those observed in cases without known germline predisposition. Specifically, the proportion of non-Ig/TCR deletions containing RSS motifs at one or both breakpoints was similar between variant carriers and non-carriers (**Figure 4D**). Multivariable negative binomial regression models adjusted for age at diagnosis and molecular subtype confirmed that the number of non-Ig/TCR deletions with at least one RSS was not significantly different between P/LP variant carriers and non-carriers (β: 0.04, 95% CI -0.19 to 0.27; p=0.73 for the lenient variant list; β: -0.18, 95 % CI -0.58 to 0.22; p=0.37 for the stringent list); We found similar results for non-Ig/TCR SVs with at least one RSS at the breakpoint [lenient variant list: β: 0.07, 95%CI: -0.12, 0.27 (p=0.47); stringent variant list: β: -0.08, 95%CI: -0.42, 0.25 (p=0.62)] (**Figure 4E**). Therefore, these data support a model in which germline predisposition variants in B-ALL contribute to the likelihood of acquiring RAG-mediated SV drivers of B-ALL due to delayed transitions in early B cell development, but once cells are transformed the burden of overall SVs in individuals with or without large-effect germline predisposition alleles appears similar.

## Discussion

The pathogenesis of B-ALL is thought to follow a multi-step model of leukemogenesis, beginning with the formation of a pre-leukemic clone, followed by the acquisition of distinct constellations of somatic structural rearrangements that drive progression toward overt malignancy (53–56). Deciphering how germline genetic variation influences this initiation process has been constrained by several experimental barriers. By systematically modeling B-ALL predisposition alleles within a tractable human *in vitro* differentiation platform, our study directly interrogates the functional effects of inherited mutations during these critical windows of human B-lymphoid development. In doing so, this methodology successfully bridges a critical knowledge gap regarding the precise temporal and cellular contexts in which germline variants exert their effects, providing a standardized framework to link inherited B-ALL risk to altered human B-lymphopoiesis. Our study provides several key advantages compared to traditional, patient-based approaches. First, our *in vitro* B-cell differentiation platform enriches for rare, transient lymphoid progenitor populations that are otherwise inaccessible and difficult to manipulate in primary human tissues, enabling direct investigation of a critical window in early B-cell development. Second, by leveraging efficient genome editing in primary human HSPCs, we establish an isogenic system to assess the functional impact of germline variants associated with B-ALL predisposition under controlled differentiation conditions. Third, although prior studies have examined individual genes, our work represents the first systematic, functionally grounded interrogation of multiple high-penetrance B-ALL predisposition alleles evaluated in parallel through an integrative and highly sensitive perturbation screen. Importantly, our study represents one of the few applications of arrayed perturbation screens in primary human cell systems. Whereas previous studies have largely focused on mature immune cells or immortalized cell lines, our platform enables systematic functional interrogation of inherited leukemia predisposition during human lymphopoiesis (57).

Through this systematic approach, we demonstrate that these diverse risk variants converge on specific developmental transition stages in early B lymphopoiesis, even when overall B-cell output remains largely preserved. Notably, the affected stages correspond to periods of elevated RAG recombinase expression and active immunoglobulin gene segment recombination, raising the possibility that prolonged residence within these vulnerable cellular states increases the opportunity for acquisition of secondary leukemogenic events, as previously suggested (48,58). The absence of shared downstream transcriptional programs further suggests that the developmental context in which these mutations act might be critical for leukemogenesis. Notably, the stage-specific differentiation blocks and impact on B cell output were gene-dependent, with *IKZF1* disruption showing the most profound effect. The apparent DN activity of one of the *IKZF1*-targeting sgRNAs further highlights the importance of modeling variant-specific effects to better delineate disease mechanisms. These findings collectively underscore the power of integrating functional genomics with developmental hematopoiesis to decode inherited leukemia risk.

While our study provides the first systematic functional dissection of high-penetrance B-ALL predisposition genes in isogenic human models, several limitations merit discussion. First, CRISPR-induced frameshift alleles in HSPCs may not fully recapitulate the nuanced effects of patient-specific missense or hypomorphic alleles. To fully resolve these patient-specific contexts, future studies incorporating precise allele-specific engineering through the use of adenine or cytosine base editors, and long-term modeling of clonal evolution will be needed (59,60). Second, although our culture system robustly models early B-cell development, it does not capture the full *in vivo* microenvironment or selection pressures that may modulate leukemogenesis in a patient context (61). Finally, while we focused on nine well-characterized predisposition genes, expanding this approach to include common risk alleles will further refine variant-to-function mapping across broader polygenic cohorts in B-ALL and allow for the discovery of novel regulators of B-cell development and B-ALL predisposition (Lee *et al*., submitted).

## Acknowledgements

We are grateful to members of the Sankaran Laboratory for their valuable feedback and helpful discussion. The lab of V.G.S. is supported by the National Institutes of Health (NIH) grants R01CA292941, R01CA265726, R01DK103794, R01HL146500, the Howard Hughes Medical Institute, the Edward P. Evans Foundation, Alex’s Lemonade Stand Foundation, Blood Cancer United, the Gates Foundation, the Leona M. and Harry B. Helmsley Charitable Trust, and philanthropic funding in memory of Jan Ellen Paradise, MD through Boston Children’s Hospital. V.G.S. is an Investigator of the Howard Hughes Medical Institute. L.W. is supported by the American Society for Hematology Scholar Award, the Damon Runyon–St. Jude Pediatric Cancer Research Fellowship Award, the Innovative Basic Science Research Award Claudia Adams Barr Program in Cancer Research (DFCI), the Translational Investigator Service Award at Boston Children’s Hospital, the Pedals for Pediatrics Research Grant, the Boston Children’s Hospital Office of Faculty Development/Basic & Clinical Translational Research Executive Committees Faculty Career Development Fellowship. A-L.N. is supported by the EMBO Postdoctoral Fellowship (ALTF 209-2024) and a Pedals for Pediatrics Research Grant. A.J.L. is supported by an NIH T32 training grant (5 T32 HL66987-22). L.M. was supported by the Boehringer Ingelheim Foundation (BIF) MD Fellowship. A.Z. is supported by an NIH MSTP grant (5 T32 GM144273-03). S.A.H. was supported by grants from the Harvard College Research Program (HCRP) and the Program for Research in Science and Engineering (PRISE). A.J.d.S. is a Scholar of Blood Cancer United.

## Author contributions

Conceptualization: L.W. and V.G.S.; Methodology: L.W., A.-L.N., A.J.L., S.B., S.M., P.M., L.M., J.P., A.Z., S.A.H.; Computational analyses: A.J.L., T.L.; Experiments: L.W., A.-L.N., A.J.L., S.B., S.M., P.M., L.M., J.P., A.Z., S.A.H.; Resources: C.W., A.C., T.F.; Visualization: L.W., A.-L.N., A.J.L., V.G.S; Supervision: V.G.S; Writing – original draft: L.W., A.-L.N., V.G.S.; Writing – review and editing: L.W., A.-L.N., A.J.L., S.B., S.M., P.M., L.M., T.L., C.W., A.C., J.P., T.F., M.A., A.Z., S.A.H., A.J.d.S., V.G.S.

## Materials and Methods

### Primary cell culture

B cell progenitors were derived from differentiation of cord blood derived human hematopoietic stem and progenitor cells (HSPCs). HSPCs were purified from discarded umbilical cord blood samples of healthy newborns using the EasySep Human CD34 Positive Selection Kit II following pre-enrichment with RosetteSep Pre-enrichment cocktail (Stem Cell Technologies) and mononuclear cell isolation on a Ficoll-Paque (GE Healthcare) density gradient. Discarded cord blood units were obtained from the Pasquarello Tissue Bank at Dana-Farber Cancer Institute (IBC-P00000180). Cells were initially cultured and expanded at 37 °C and 5% CO2 in serum-free medium consisting of StemSpan II medium (Stem Cell Technologies) supplemented with CC100 cytokine cocktail (Stem Cell Technologies) and 50 ng/ml TPO (Peprotech) (62). Confluency was maintained between 5 × 10^5 and 1 × 10^6 cells per ml. After 5 days of expansion culture, HSPCs were co-cultured with the MS-5 stromal layer to facilitate B cell differentiation. Cells were maintained for 3-5 weeks with bi-weekly feedings with IMDM (StemCell) containing 5% FBS, 50 µM 2-Mercaptoethanol 1% penicillin/streptomycin and 20 ng/mL recombinant IL-7 (StemCell) (63). Naive B cells were isolated from discarded cord blood units using the EasySep Human Naive B cell isolation kit according to the manufacturer’s instructions (Stem Cell technologies).

### Cell lines

MS-5 cells (DSMZ) were cultured at 37 °C in IMDM (StemCell) supplemented with 10% FBS, 50 uM 2-Mercaptoethanol (Gibco), 1% L-Glutamine and 1% penicillin/streptomycin. 24-48 hours prior to establishment of co-culture with HSPCs, cells were pre-plated in a 24 well plate at 3 × 10^4 per well.

### Morphological analysis of primary cell cultures

To analyze the morphology, CD19^+^ cells were purified from differentiating primary cell cultures or peripheral mononuclear cells using anti-CD19 MicroBeads (Miltenyi) according to the manufacturer’s instructions. Subsequently, 70-100,000 cells were resuspended in 2% FBS in PBS and centrifuged using a Cytospin 4 centrifuge (Thermo Scientific) at 300 g for 4 minutes with low acceleration. Air-dried slides were stained using May-Grünwald solution (Sigma Aldrich, MG1L) for 5 minutes, rinsed 4 times for 30 seconds in water, and stained using Giemsa solution (Sigma Aldrich, 32884) for 15 minutes. Slides were rinsed 6 times for 30 seconds with water and mounted using a vectashield mounting medium (Vector Laboratories). Slides were examined using an Axio Imager Z2 microscope (Zeiss) and images were analyzed using ImageJ.

### Real-time quantitative PCR (RT-qPCR)

Total RNA was extracted using the AllPrep DNA/RNA Mini Kit (QIAGEN, 80204). 100-500 ng of RNA was reverse transcribed using the iSCRIPT cDNA Synthesis kit (Bio-Rad, 1708891) following the manufacturer’s instructions. RT-qPCR was performed using iQ SYBR Green Supermix (Biorad, 1708880) and the CFX384 Touch Real-Time PCR Detection System (Bio-Rad). Primers used are listed in **Supplementary Table 1**.

### CRISPR/Cas9-genome editing and analysis

CRISPR/Cas9 genome editing of HSPCs was performed while cells were in progenitor maintenance media (62). Electroporation was performed 48 hours after thawing of cord-blood derived CD34^+^ HSPCs using the Lonza 4D Nucleofector kit. The RNP complex was made by combining 100 pmol Cas9 (IDT) and 100 pmol modified sgRNA (Synthego) targeting *AAVS1, CDKN2A, ETV6, IKZF1, NBN, PAX5, SH2B3, TCF3, TP53* or *USP9X* using single synthetic guide RNAs (**Supplementary Table 2**) to introduce indels and incubating at room temperature for 30 min. Between 1 × 10^5 and 5 × 10^5 HSPCs were resuspended in 20 µl P3 solution with 1 uL electroporation enhancer (IDT) and mixed with the RNP to undergo nucleofection with program DZ-100 (HSPCs). Cells were returned to HSC medium and editing efficiency was measured at 48-72 hours after electroporation, unless otherwise indicated.

### Flow cytometry

Cells were harvested, filtered, washed with PBS supplemented with 2% FBS, incubated with FcR Blocking Reagent (Miltenyi) and stained with the following panel of antibodies: anti-CD34-Alexa488 (1:50, BioLegend; RRID: AB_1937204), anti-CD10-PE (1:50, Beckman-Coulter, AB_131294), anti-CD19-BV421 (1:50, BD, AB_11153299), anti-CD20-Pe-Cy7 (1:50 BD, AB_1727450), anti-CD33-APC (1:50, BioLegend, AB_314351) or anti-CD45-APC (1:50, Thermo Fisher Scientific, AB_10667894) for 30 minutes at 4°C. Cells were stained for viability using 7-AAD (1:300, BioLegend). Flow cytometric analyses were conducted on a BD Fortessa analyzer and all data were analyzed using FlowJo software v10.9.0.

### B cell receptor sequencing

B cell receptor (BCR) sequencing was performed using the NEBNext Immune Sequencing Kit according to the manufacturer’s instructions. In brief, HSPCs from six independent donors were differentiated using the *in vitro* platform and compared to *bona fide* Naive B cells isolated from a healthy donor. The resulting libraries were sequenced using the Illumina Miseq 600 cycles kit. Bulk B Cell Repertoire Analysis was performed using the Immcantation Framework and associated workflows (64,65).

### Editing efficiency

Genomic DNA (gDNA) was extracted from edited cells using the following kits, depending on the number of input cells (QIAamp DNA Micro Kit: 56304, QIAamp DNA Mini Kit: 51304 Qiagen DNeasy Blood & Tissue Kit: 69504 and NORGEN genomic DNA isolation Kit: 24700). NCBI’s PrimerBLAST and Benchling was used to design primers specific to the target locus for an amplicon of 300-500 bp of size. PCR amplification was performed on 20 - 100ng of gDNA using Platinum II Hot-Start PCR Master Mix 2x (Thermo Fisher Scientific) or Q5 Hot Start High-Fidelity 2X Master Mix (New England Biolabs) with specific primers (**Supplementary Table 2**). PCR products were subsequently purified with the PCR & DNA Cleanup kit (New England Biolabs) and subjected to Sanger sequencing. Editing efficiency was analyzed using the Inference of CRISPR Edits (ICE) (ice.synthego.com) tool.

### Single-cell RNA sequencing

Droplet-based digital 3’-end single cell RNA sequencing (scRNA-seq) was performed on a Chromium Single-Cell Controller (10X Genomics) using the Chromium Next GEM Single Cell 3’ Reagent Kit v3.1 according to the manufacturer’s instructions. Cells were processed according to the TotalSeq™ -A Antibodies (**Supplementary Table 3**) and Cell Hashing with 10x Single Cell 3’ Reagent Kit v3.1 (Dual Index) Protocol (Biolegend and 10x Genomics). Differentiating cells were harvested from MS-5 stroma layers using PBS after 21 days of co-culture. Prior to FACS enrichment, cells were conjugated with Hash Tag Oligonucleotides at 1uL per 1 × 10^6 cells. Cells were stained on ice for 30 min with the following antibodies: CD10-PE, CD19-BV421, CD45-APC and CD34-Alexa Fluor 488 (all 1:50). Cell viability was assessed using 7-AAD (1:300). Cells were washed and sorted by FACS for viability and expression of human CD45 to remove any remaining MS-5 cells from the culture. Cells were sorted into 5-mL 0.1% BSA in PBS-coated tubes and were spun down at 400 x g for 5 minutes at 4°C. The entire supernatant was carefully removed and 1mL of PBS+0.05% BSA was added. Cells were then counted using a hemocytometer. Cell viability as determined by Trypan Blue was >95%. Cells were washed with 5mL PBS+0.05% BSA and resuspended at a final concentration of 1,000 cells / μL and kept on ice. Single-cells were immediately processed using v3.1 Chemistry Dual Index kits, using 20μL of cell suspension and 23.2 μL of water on the cell suspension loading step. Libraries were sequenced using a 28 base pair (Read 1), 10 base pair (Index 1), 10 base pair (Index 2), 90 base pair (Read 2) configuration on Novaseq S3 instruments.

### Single-cell RNA sequencing data analysis

Demultiplexed FASTQ files were processed with Cell Ranger (10x Genomics) for both gene expression and hashtag oligonucleotide (HTO) libraries using the GRCh38-2024-A reference. Cells were assigned to their sample of origin by hashtag demultiplexing using the HTODemux. Using a standard Seurat pipeline (v4.4.0), aligned scRNA-seq reads from each sample were aggregated into a single object. Cells with fewer than 200 or more than 5,000 detected genes, or a mitochondrial unique molecular identifier (UMI) fraction higher than 20% were removed. Potential doublets were identified and excluded using ScrubletR (v0.2.0) with a doublet score threshold of >0.4. Cell identities were assigned by projecting the cells from each sample individually onto a human B cell development reference (https://github.com/andygxzeng/b_development_map) and bone marrow reference (https://github.com/andygxzeng/BoneMarrowMap) using Symphony, followed by minor curation of cell type annotations to align them with cell types identified in the *in vitro* differentiation culture (28,42,66). To examine the global transcriptional similarity between *in vitro*-derived cells and bona fide differentiating B cells, HSPC-derived cells from week 3 and 5 were co-embedded with native B cells from orthogonal datasets onto the human B-cell development reference (41). The integrated cells were partitioned into 100 subclusters using k-means clustering. After excluding subclusters containing fewer than 50 cells, Pearson correlation coefficients between *in vitro*- and *in vivo*-derived subclusters were calculated based on quantile-normalized, log2-transformed pseudobulk expression matrices generated from genes expressed in more than 10% of cells. Gene set enrichment analysis was performed using the fgsea package (https://github.com/ctlab/fgsea/) using the normalized pseudobulk expression data with gene sets from the MSigDB Hallmark and GO Biological Process collections. Transcriptional signature scores were assessed by using Seurat’s AddModuleScore function (28,42).

### Germline P/LP variant calling in B-ALL patient cohort

To investigate the frequency of RAG recombination-mediated structural variants (SVs) in B-ALL patients harboring germline pathogenic/likely pathogenic variants in predisposition genes and in non-carrier patients, we leveraged available sequencing data in a cohort of 1,491 patients enrolled in Children’s Oncology Group (COG) trials from the Molecular Profiling to Predict Responses to Therapy (MP2PRT) study (dbGaP accession number phs002005.v1.p1) (51). Raw germline sequencing data and somatic SV calls were available for n=1,491 B-ALL patients via the National Cancer Institute Genomics Data Commons (GDC) Data Portal through an approved dbGaP request. We used bamSliceR (67) to download specific genomic regions from the patient BAM files, encompassing the following genes of interest: *IKZF1* (sliced region, chr7:50304716-50405101), *NBN* (chr8:89933331-89984667), *CDKN2A* (chr9:21967752-21974857), *PAX5* (chr9:36833269-37034268), *ETV6* (chr12:11649674-11895377), SH2B3 (chr12:111405923-111451623), *TP53* (chr17:7668421-7687490), *TCF3* (chr19:1609292-1652615), *RUNX1* (chr21:34787801-35049302), *USP9X* (chrX:41085445-41236579). Variant calling and filtering were performed using GATK HaplotypeCaller and VariantFiltration (68). Variants that passed QC (QD > 2, FS > 60, MQ < 40, and SOR > 3) and had < 0.1% population frequency were retained and annotated using Annovar (69), PeCanPIE (70), Ensembl Variant Effect Predictor (VEP) (71), AlphaMissense (72), and ClinVar. Variants predicted to be “HIGH” impact by VEP and/or predicted to be P/LP by ClinVar were included in a “stringent” list of P/LP variants. Additional variants predicted to be “MODERATE” impact by VEP and to be “likely pathogenic” by AlphaMissense, or variants in *IKZF1* predicted to be P/LP in prior functional analyses (12), were included in a larger “lenient” list of P/LP variants (73).

### RAG-mediated SVs in P/LP variant carriers

SV calls generated in MP2PRT B-ALL patients using the BRASS SV calling pipeline were downloaded from the NCI GDC Data Portal. We predicted whether SVs were likely to have been formed by RAG recombination (i.e., RAG-mediated SVs) as previously described (52,74). In brief, we determined the presence of full-length RSS motifs in the sequences flanking +/-50bp of each SV breakpoint using the Find Individual Motif Occurrences (FIMO) tool in MEME suite v5.5.5 (75,76). We further annotated RAG-mediated SVs as “on-target” or “off-target” according to whether at least one breakpoint overlapped with Ig/TCR gene regions or whether both breakpoints were +/-1000 bp outside of Ig/TCR regions, respectively. We compared the total number of different SV types, the number of RAG-mediated SVs and deletions, and the fraction of total SVs/deletions that were RAG-mediated between P/LP variant carriers and non-carriers among n=1,491 MP2PRT B-ALL patients using the Wilcoxon rank-sum test.

## Data availability

All raw and processed data generated in this study have been deposited in the Gene Expression Omnibus (GEO) repository and will be available at the time of publication. Joint single-cell multi-omic data of human bone marrow used for orthogonal validation were obtained from the GEO repository (GSE219015 and GSE194122). No original code was generated in this study.

**Figure S1:**
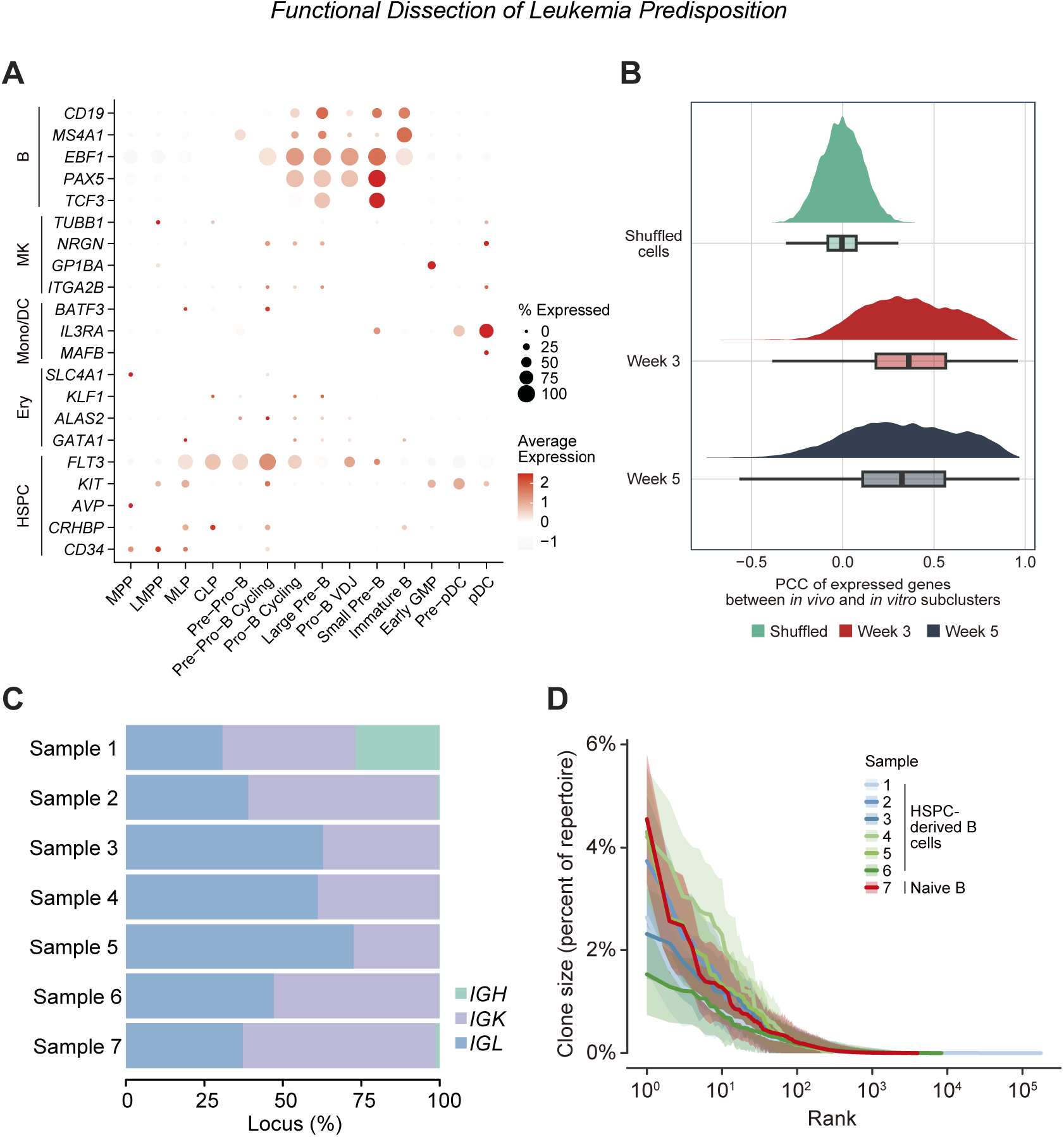
Molecular characterization of HSPC-derived B cells. **(A)** Dot plot showing the expression of marker genes for HSPCs and the erythroid (Ery), monocyte and dendritic cells (Mono/DC), megakaryocyte (MK), and B cell lineages across stages of B cell development. **(B)** Ridge plot displaying the Pearson Correlation Coefficient (PCC) of RNA expression between HSPC-derived B cells at week 3 and 5 of *in vitro* differentiation and a single-cell reference map of B cell differentiation in vivo, across all expressed genes. **(C)** Quantification of mapped V(D)J rearrangements originating from the immunoglobulin heavy (IGH), kappa light (IGK), and lambda light (IGL) chain loci across HSPC-derived B cells (*n*=6; samples 1-6) and naive B cells (*n*=1; sample 7). **(D)** Clonal abundance distributions of BCR repertoires from HSPC-derived B cells (*n*=6) and PBMC-derived B cells (*n*=1).

**Figure S2:**
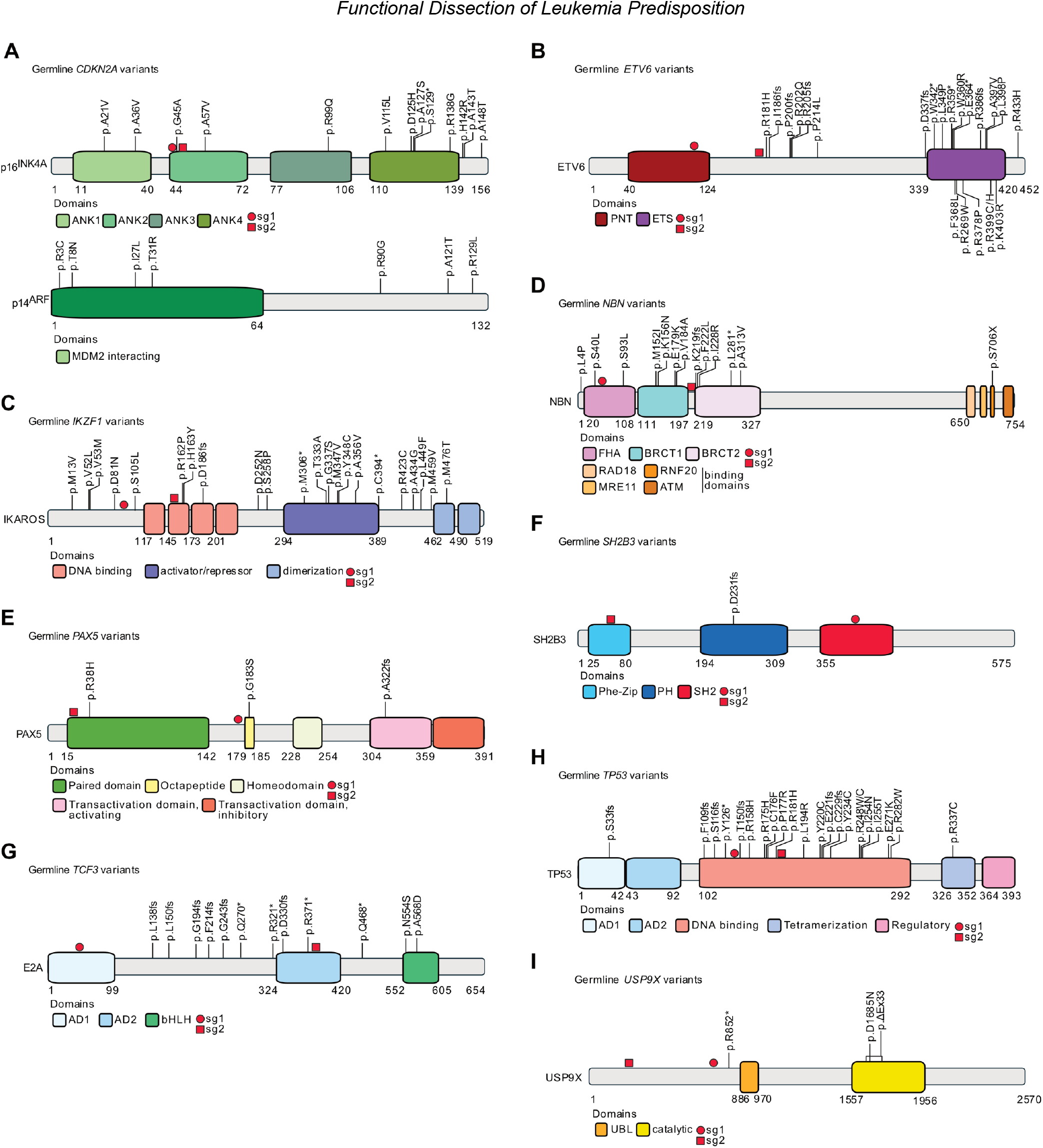
Location of B-ALL risk variants and sgRNAs. **(A-I)** Location of previously reported deleterious germline B-ALL risk variants mapped onto the proteins encoded by **(A)** *CDKN2A*, **(B)** *ETV6*, **(C)** *IKZF1*, **(D)** *NBN*, **(E)** *PAX5*, **(F)** *SH2B3*, **(G)** *TCF3*, **(H)** *TP53* and **(I)** *USP9X* (12–19,35). Protein domains are indicated by color and location of sgRNAs are indicated by a red circle (sgRNA_1) or red square (sgRNA_2), respectively. ANK: ankyrin repeat; PNT: pointed; FHA: forkhead-associated; BRCT: BRCA1 C-Terminus; Phe-Zip: Phenylalanine zipper; PH: Pleckstrin homology; SH2: Src Homology 2; AD: Activation domain; bLHL: basic helix-loop-helix; UBL: ubiquitin-like.

**Figure S3:**
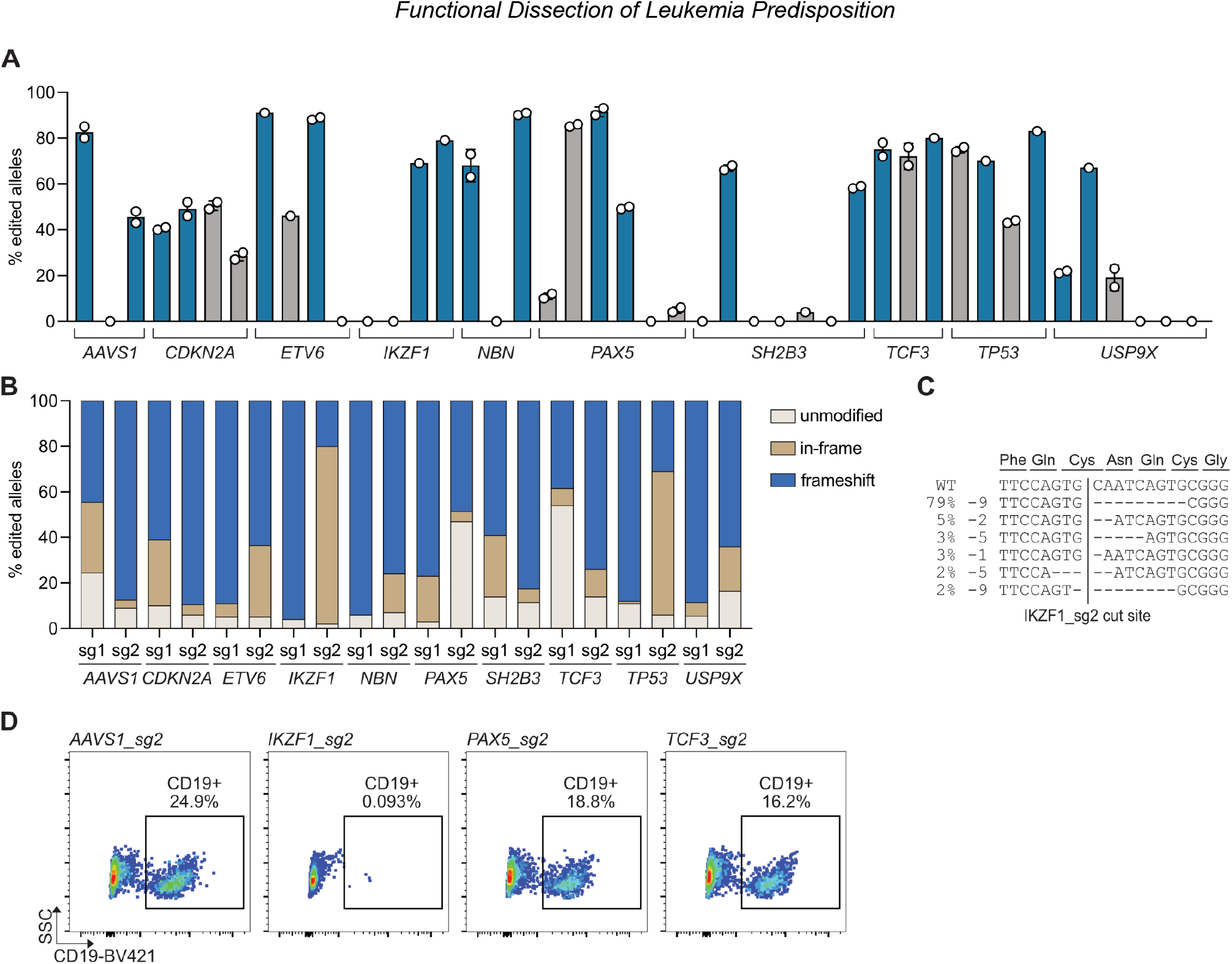
Optimization of CRISPR/Cas9 editing in primary CD34^+^ HSPCs. **(A)** Screening of sgRNAs for editing efficiency across B-ALL predisposition loci at day 5 post nucleofection of primary CD34^+^ HSPCs. Blue bars represent sgRNAs that were subsequently selected for the arrayed perturbation screen. **(B)** Distribution of editing outcomes for selected guide pairs per target. **(C)** Characterization of editing outcomes generated by IKZF1_sg2 aligned to the reference (WT) sequence. **(D)** Representative flow cytometric analysis of B-cell differentiation following CRISPR/Cas9 editing with indicated sgRNAs.

**Figure S4:**
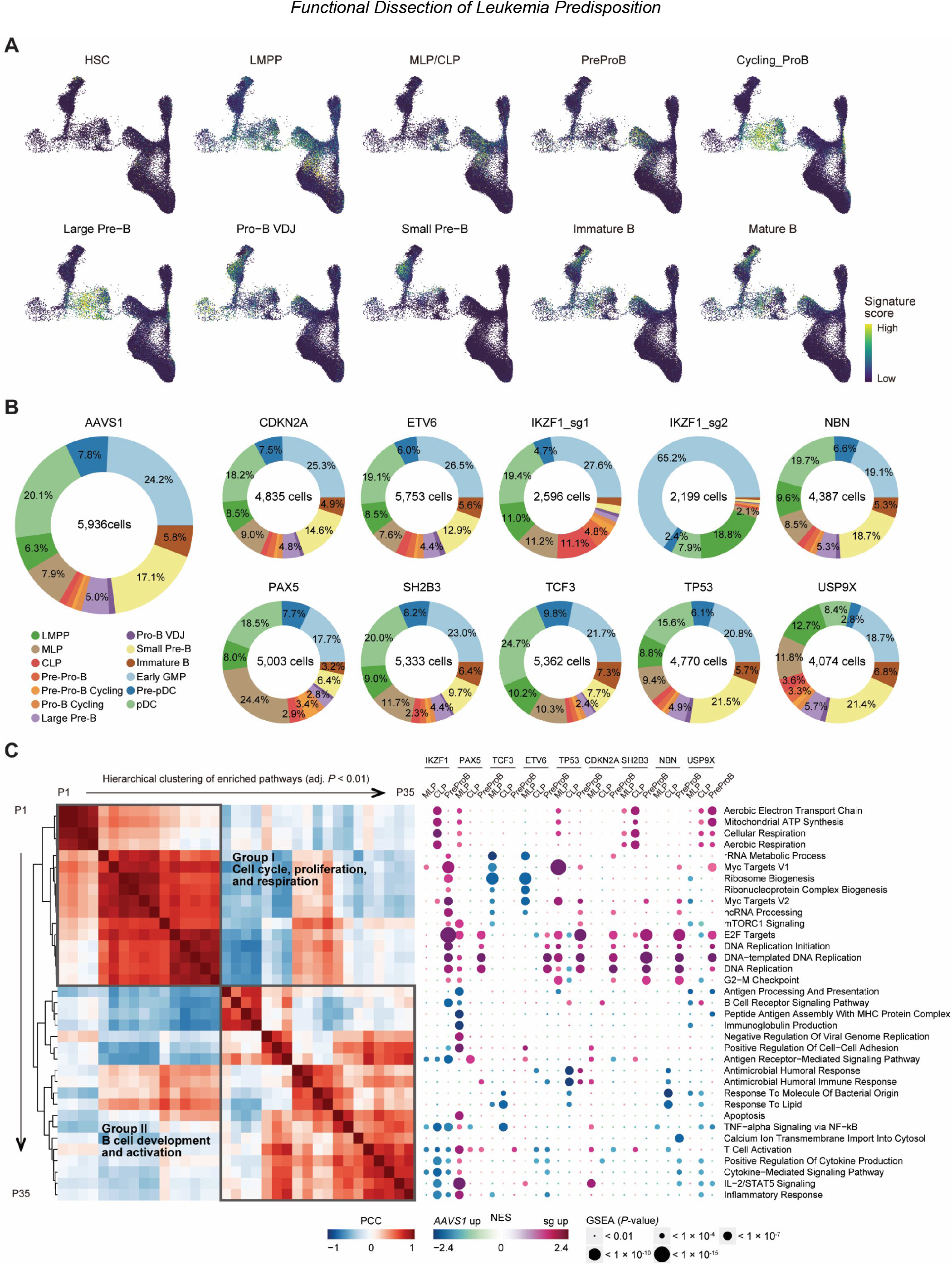
ScRNA-seq analysis of B-ALL predisposition alleles. **(A)** UMAP showing signature scores of top five marker genes identified from a single-cell reference map of hematopoietic differentiation for each cell type within B-cell development. **(B)** Pie charts showing the proportion of cells for each sgRNA-targeted perturbation and the corresponding number of cells captured. **(C)** Gene set enrichment analysis (GSEA) showing significantly enriched pathways (FDR < 0.01) in each sgRNA-mediated perturbation compared with *AAVS1* controls, based on pseudobulk counts within stalled states of B cell differentiation, using Hallmark and GO Biological Processes gene sets. The enriched pathways are organized by hierarchical clustering based on normalized enrichment score (NES) similarity.

